# Investigating the influence of experimentally induced central sensitisation on pain prediction error encoding in healthy individuals: a novel virtual reality protocol

**DOI:** 10.1101/2025.01.30.635687

**Authors:** Federico Palmisani, Jawahar Sri Prakash Thiyagarajan, Joe Horsey, Sam Hughes, Sonia Medina

## Abstract

Persistent mismatches between predicted and actual pain-related signals, namely prediction errors (PEs), can cause maladaptive overestimation of pain intensity, a common feature of chronic pain states. Experimental protocols used to assess the contribution of central sensitisation (CS) to dysregulated prediction systems are lacking. To address this, we implemented a novel virtual reality (VR) paradigm to evoke PEs during mechanical stimulation following experimentally induced CS via the high-frequency stimulation (HFS) model. Twenty healthy volunteers underwent HFS on the right forearm. Mechanical pain sensitivity (MPS) was assessed through pinprick stimuli before and 30 minutes post-HFS to evaluate secondary hyperalgesia. Following this, participants received mechanical stimuli at proximal (sensitised area) and distal (non-sensitised area) points from the HFS site, with visual cues presented on their arm via VR alongside hand tracking technology indicating the stimulus location, allowing participants to make pain predictions. Cues were either congruent (matching) or incongruent (mismatching) with the actual stimulus site, to evoke PEs. Results showed that MPS significantly increased following HFS, confirming secondary hyperalgesia. Stimuli in sensitised areas induced more pain than in non-sensitised areas. Incongruent cues successfully elicited PEs across all locations, however, expectations modulated pain perception only in non-sensitised areas. Similarly, during incongruent trials, PEs diminished over time (reflecting adaptive learning) only in non-sensitised areas. These data demonstrate that pain expectations can influence pain perception differently in centrally sensitised and non-sensitised states. We propose this protocol as a good candidate to assess how cognitive and psychological manipulations influence PEs at various stages of CS.

## Introduction

The predictive coding model of pain suggests that the brain continuously anticipates sensory input to prepare for adaptive responses [57; 60]. Predictions are shaped by prior learning and expectations [12; 34]. When the predicted pain intensity mismatches pain perception in the presence of a given sensory input, prediction errors (PEs) arise [13]. PEs allow to adjust future predictions, dynamically modulating incoming signals to fine-tune pain responses based on whether it has undefrestimated or overestimated the incoming threat, aiding protective reflexes [60]. However, during chronic pain, individuals may begin to overestimate the danger posed by sensory stimuli due to fear of pain [73] and pain catastrophising [23; 82]. On the other hand, persistent afferent signal amplification at the dorsal horn, known as central sensitisation (CS), can lead to heightened sensitivity to both noxious and non-noxious mechanical stimuli [4; 32], further biasing the sense of immediate threat [25; 62] and making the brain overly sensitive to pain, even in the absence of nociceptive input.

Understanding the specific role of CS in driving persistent PEs, where the brain fails to adjust its pain predictions despite changes in sensory input, is therefore crucial for designing personalised treatments [31], that target the underlying root of maladaptive pain learning [3; 37]. Nevertheless, experimental protocols able to assess this are currently lacking. The high frequency stimulation (HFS) model can induce prolonged secondary hyperalgesia in healthy individuals within a well-defined heterotopic area [17; 42; 70], a phenomenon attributed to CS. Considering this, we introduce a novel approach where we can induce PEs following HFS, by presenting visual cues that indicate the location of upcoming stimuli, either inside or outside of the sensitised area, and manipulating the congruency between the cue and the actual stimulation site, that gives rise to PEs. By integrating virtual reality (VR) with hand-tracking technology [14], we can deliver visual cues with a high level of experimental control [39] while inducing high levels of embodiment [53], as it allows participants to continuously view a virtual representation of their arm while remaining unaware of the actual location of the stimuli in relation to the cued site. We proposed that this strategy can allow us to test how the presence of PEs interacts with the extent of secondary hyperalgesia to result in different levels of perceived pain.

In this study, we explored pain modulatory responses due to experimentally induced PEs following HFS using VR in a healthy cohort. Our preregistered hypotheses [46] were: i) HFS would increase mechanical pain sensitivity measures; ii) PEs would influence pain perception: if predicted pain at the attended location is lower than perceived pain, the result would be lower pain perception compared to a match between predicted and perceived pain. If predicted pain is higher than perceived pain, the result would be higher pain perception compared to a match between predicted and perceived pain. Crucially, we set out to explore how these bidirectional relationships vary depending on whether stimulation is delivered in a sensitised versus non-sensitised area.

## Methods

All methods and data analyses adhered to the preregistered study plan available on OSF (see Palmisani et al. [46]).

### Participants

A total of 20 healthy individuals, aged between 20 and 31 (mean age: 25, SD = 2, 15 males), participated in this study. The sample included individuals aged over 18 years identifying as healthy. Exclusion criteria included the presence of chronic illnesses or current pain, a history of epilepsy, substance or alcohol abuse, skin conditions such as eczema, and any ongoing psychological conditions requiring psychoactive medications, unless the dosage had remained stable for at least three months. Additionally, participants were excluded if they consumed more than eight caffeinated beverages or smoked more than five cigarettes per day. Participants were instructed to abstain from alcohol, nicotine, painkillers, and all but a single caffeinated drink on the day of the study. Each participant provided written informed consent prior to participating in the study. The experiment was approved by the Health Research Authority and Health and Care Research Wales Ethics Committee (reference: 22/HRA/4672).

### Experimental Design

This study employed a repeated-measures, within-subject design to investigate the influence of top-down predictive processes on pain perception during peripherally induced central sensitisation. Each participant completed a single session in the laboratory. Compliance with study lifestyle guidelines was assessed at the beginning of the session. In order to ensure the absence of any ongoing potentially confounding pain, participants also answered the question ‘Are you in any pain today’ with a numerical score on a numerical rating scale (NRS) that ranged from 0 (‘no pain at all’) to 100 (‘worst pain imaginable’). All participants underwent study procedures in the same order.

### High-frequency stimulation (HFS) model

Electrical stimulation was applied to the centre of the right volar forearm using an epicutaneous pin electrode. This electrode consisted of a circular array of 15 cathodal pins (individual pin diameter: 0.2 mm; length: 1 mm; total array diameter: 10 mm; area: 79 mm^2^), surrounded by a stainless-steel anode (inner diameter: 20 mm; outer diameter: 40 mm). A constant current stimulator (pulse width: 2 ms; DS7, Digitimer Ltd, Welwyn Garden City, UK) delivered the stimuli. Before electrode placement, participants’ forearms were cleaned with isopropyl alcohol. The stimulation site was marked using a circular template matching the electrode’s size, with one reference points spaced 1 cm apart along eight radial axes. A corresponding grid was drawn on the left volar forearm to ensure consistency during control assessments. Individual electrical detection thresholds (EDTs) were determined using the method of limits [21]. Stimuli were first delivered at an intensity of 0.05 mA, increasing in steps of 0.05 mA until participants reported a distinct sensation. The intensity was then reduced in 0.01 mA increments until the sensation disappeared and subsequently increased again in 0.01 mA increments until the sensation returned. This procedure was repeated three times, and the average of six measurements (three ascending and three descending) was calculated to establish the EDT for each participant. Immediately after calculating participants’ EDT, HFS was administered using a pulse generator (D4030; Digitimer Ltd). The protocol consisted of five trains of 100 Hz stimuli, each lasting one second, with ten-second intervals between trains. The stimulation intensity was set at 20 times the individual EDT. Following each train, participants rated their perceived pain intensity in an NRS.

### Mechanical pain sensitivity (MPS) assessments

Identical MPS assessments were carried out in both forearms at baseline and 30 minutes post-HFS using a modified version of the DFNS MPS protocol [77]. Three pinprick stimuli with forces of 128 mN, 256 mN, and 512 mN were applied five times each in a pseudo-randomized sequence within a 1 cm area surrounding the electrode site. To minimize windup effects, the locations of the stimuli were alternated across four quadrants, using the landmarks previously drawn as a guide. After each stimulus, participants rated their perceived pain intensity on an NRS. To control for potential order effects, the sequence in which each forearm was tested (left or right) was counterbalanced across participants but kept consistent within participants.

### Procedure

#### Virtual Reality (VR) Setup

Following 30 minutes post-HFS MPS assessments, participants were introduced to the VR environment. Seated comfortably with their right arm positioned on a pillow placed on a table in front of them, participants wore an HTC VIVE 2 PRO headset, equipped with an Ultraleap hand-tracking camera to display a real-time virtual representation of their arm. The virtual arm included a circle on the location consistent with the HFS electrode placement. Participants were asked to direct their gaze towards their right arm and move their fingers to ensure the virtual representation of their arm responded accurately to their real movements and induce a sense of embodiment of the virtual arm. A total of eight locations were selected for stimulation during the VR task, including four locations on the heterotopic area proximally to the electrode across axes 1, 3, 5 and 7 (from the eight axes originally drawn for MPS assessment) and four distal locations (3cm away from proximal locations) on the same axes (Figure 1). All stimuli were delivered with a 512 mN pinprick, which represents a deviation from the original protocol; while our original protocol included 128 mN intensity stimuli, preliminary pilot testing indicated that participants did not report sufficient levels of pain for meaningful data analysis. Consequently, we opted to use the 512 mN pinprick stimulus, which elicited more robust pain ratings. The experimental task included three blocks of trials in total:

**Figure 1.**
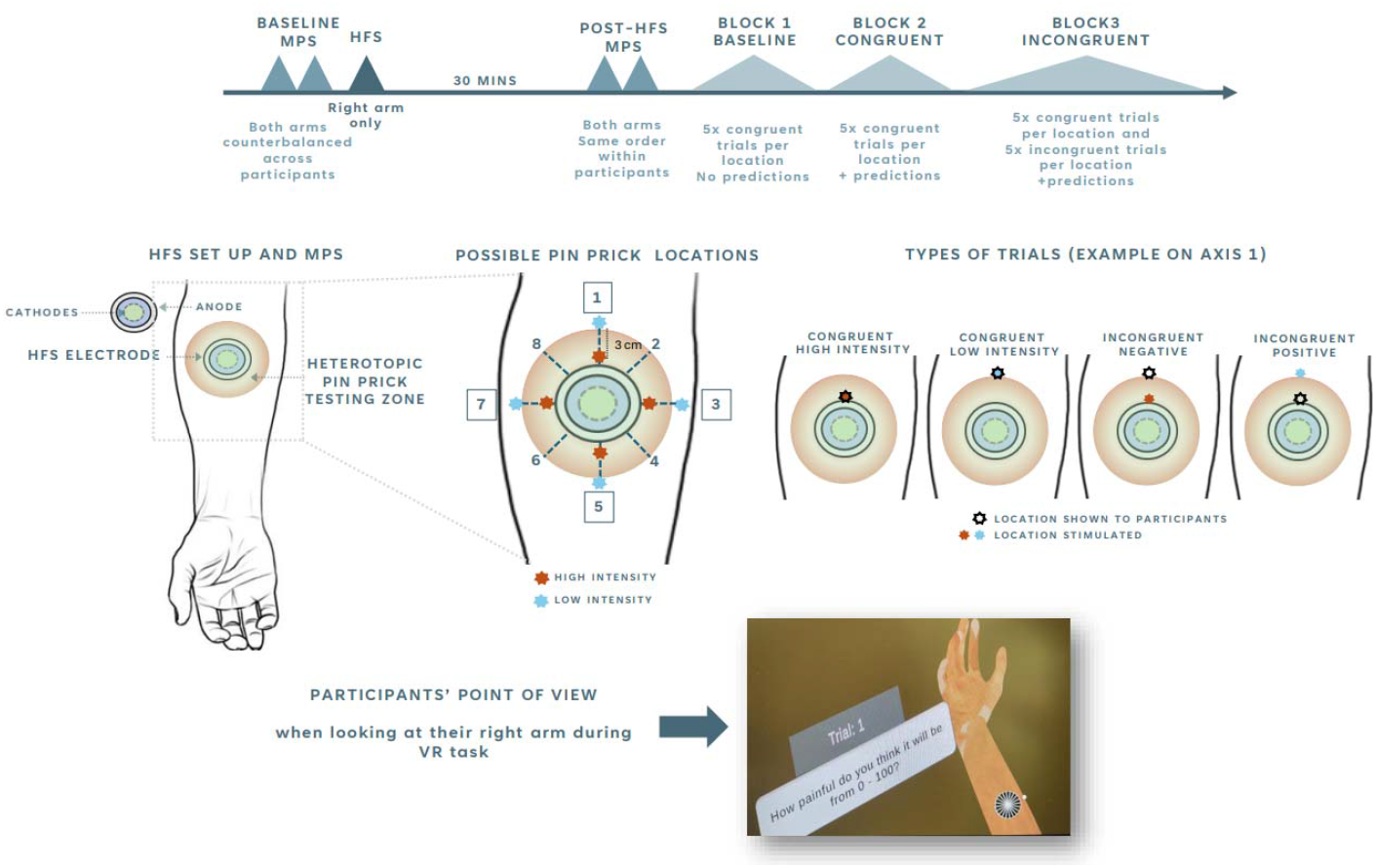
Experimental design (top panel) HFS set up and trial classification (middle panel) and VR set up (bottom panel). All participants attended a single visit and completed an identical protocol. The total duration of the visit was approximately 1.5 hours. MPS was defined as the geometric mean of 15 pinprick stimulation (5 x 128 mN, 5 x 256 mN, 5 x 512 mN). Stimuli during the VR task were delivered with a 512 mN pinprick. MPS = mechanical pain sensitivity; HFS = high-frequency stimulation. HFS set up figure adapted from Medina and Hughes (2024).

*Block 1 (baseline):* in each trial, a white dot appeared on a location of the virtual arm consistent with the location of the forthcoming stimulus (Figure 1). Approximately two second later, Experimenter 1 poked the congruent landmark location on the participant’s real right arm. Immediately after, Experimenter 2 displayed a question saying, ‘How painful was the stimulus from 0 - 100?’ next to participants right arm in the virtual environment. Participants answered the question verbally with an NRS score and Experimenter 2 triggered the next trial, where a different dot would appear on the virtual arm. There was a total of 40 consecutive trials (5 trials per stimulus location), presented in randomised order (same order across participants).

*Block 2 (congruent trials):* trials were similar to those in Block 1, however, an additional prompt was displayed next to participants’ right arms in the virtual environment. This prompt read, “How painful do you think it will be from 0 - 100?” and appeared immediately after the presentation of the forthcoming stimulus landmark. ‘How painful do you think it will be?’ immediately after the display of the forthcoming stimulus landmark. Participants provided their predicted pain intensity on an NRS. Approximately two seconds later, Experimenter 2 poked the congruent landmark location on the participant’s real right arm. The remainder of the trial sequence followed the same procedure as in Block 1. A total of 40 consecutive trials (5 trials per stimulus location) were presented, maintaining the same order as in Block 1.

*Block 3 (congruent + incongruent trials):* here, the trial sequence was identical to the one in Block 2, however, the landmark shown on the virtual arm was congruent with the location poked in 50% of the trials. These were referred to as ‘congruent trials’. The remaining trials were referred to as ‘incongruent trials’. Incongruency could be presented in two ways: i) ‘incongruent negative’ (also referred to as ‘-High’), where participants were shown a low intensity landmark but were poked in a high intensity landmark within the same axis, resulting in an outcome expected to be more negative than predicted; ii) ‘incongruent positive’ (also referred to as ‘+ Low’) where participants were shown a high intensity landmark but were poked in a low intensity landmark within the same axis, resulting in an outcome expected to be more positive than predicted. A total of 80 trials were presented in randomised order (kept consistent across participants), including 40 congruent trials and 40 incongruent trials (20 incongruent negative and 20 incongruent positive). In order to avoid additional sensitisation from temporal summation of pain, no more than to consecutive trials were performed in the same location.

Participants were instructed to maintain their gaze towards their right arm for the whole duration of the block. Between blocks, participants were given the option to take short breaks to minimise discomfort from prolonged use of the VR headset. Experimenters 1 and 2 maintained the same roles throughout the study for experimental consistency.

### Data Analysis

Analyses were performed using via MS Excel and custom-made Matlab scripts. All statistical tests were considered significant with alpha = 0.05.

#### HFS effects on MPS measures

MPS indices were defined as the geometric mean of the 15 pain ratings of the pinprick stimuli (128 mN, 256 mN and 512 mN, x5 times each). MPS changes following HFS were examined via three different strategies; first, raw MPS measures were compared within each arm via paired-samples t-tests. Secondly, we calculated standard MPS changes from baseline as follows: MPSindivudual−MPSbaseline)/SDbaseline [40; 41]. Thirdly, we computed the percentage of change from baseline MPS to post-HFS assessment. One-sample t-tests were performed for standard MPS change and percentage of change scores within each arm to assess significant group changes from 0.

#### Differences in perceived pain between high and low intensity stimuli

In order to examine whether stimuli proximal to the HFS electrode (from now on referred to as ‘high intensity’ stimuli) are perceived as more painful than stimuli located distally (from now on referred to as ‘low intensity’ stimuli), average NRS scores on perceived pain following stimuli across all 4 axes were averaged separately for high and low intensity stimuli and compared via a paired-samples t-test for Block 1, Block 2, Block 3 congruent trials and Block 3 incongruent trials. Identical analyses were performed within each separate axis.

#### Prediction accuracy

Prediction errors (PEs) were defined as delta scores between NRS predictions and following perceived pain NRS scores for each trial across four groups, i.e., Block 2, Block 3 congruent trials, and Block 3 incongruent trials (PE = NRS_perceived_ – NRS_predicted_). Resulting positive PEs indicated that participants perceived more pain than predicted, and viceversa. One-sample t-tests were performed across PEs within high and low intensity stimuli in each group. PEs significantly different from zero denoted low prediction accuracy, and viceversa.

PE differences across groups was assessed via a two-way repeated measures analysis of variance (ANOVA) with group (Block 2, Block 3 congruent, Block 3 incongruent) and stimulus intensity (high, low) as a within-subject factors.

#### Relationship between predicted and perceived pain

To test whether congruency of visual cues impacted pain perception, we compared perceived pain NRS scores between Block 2 and incongruent trials in Block 3 (separately for high and low intensity trials) by means of a paired-samples t-test. In order to explore how the extent of these pain changes related to prediction accuracy, Pearson’s correlations between PE measures and delta scores calculated from NRS from congruent trials in Block 2 minus incongruent trials in Block 3 were computed, separately for high and low intensity stimuli.

#### Exploratory analysis: examination of learning effects

Two strategies were adopted to explore whether PEs were updated with time in Block 3; firstly, PEs from the first and second halves of the trials were averaged and compared via a paired samples t-test, separately for congruent, incongruent negative and incongruent positive trials. Secondly, three separate linear regression analyses were performed for averaged PEs across participants in congruent, incongruent negative and incongruent positive trials, with PEs as dependent variable and trial number as independent variable.

## Results

### Effect of HFS in mechanical pain measures

Paired-samples t-test indicated that MPS significantly increased post-HFS within participants in the test arm (mean MPS_baseline_(SD) = 5.65(4.26), mean MPS_post-HFS_(SD) = 15.59(11.67), t_(19)_ = 5.09 *p*<0.001). One-sample t-tests for standard MPS changes from baseline within each arm revealed a significant group increase in MPS measures for the test arm (t_(19)_ = 5.09 *p*<0.001), averaging 2.33 standard deviations (Figure 2), and a significant 240% increase in MPS measures (t_(19)_ = 4.45 *p*<0.001). There were no significant changes across any of the MPS indices for measures from the control arm.

**Figure 2.**
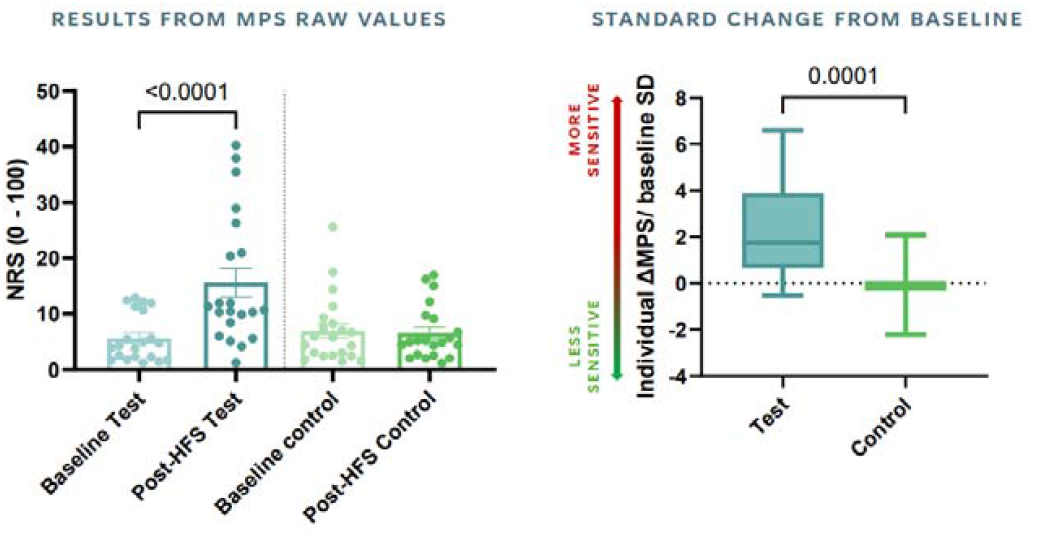
HFS effects on MPS measures. Left panel depicts differences between baseline and post-HFS assessments in raw MPS values for the test arm (blue) and the control arm (green). Error bars represent the standard error of measurement (SEM). Boxplots (right panel) represent standard changes in MPS measures from baseline for the test arm (blue) and the control arm (green). Standard MPS changes were significantly greater than zero in the test arm only, indicating the presence of secondary hyperalgesia in a constricted area surrounding the HFS site. MPS = mechanical pain sensitivity; NRS = numerical rating scale (ranged from 0 ‘no pain at all’ to 100 ‘worst pain imaginable’).

### Differences in perceived pain between high and low intensity stimuli

Paired-samples t-test revealed that stimuli located proximally to the HFS electrode (high intensity stimuli) were perceived on average as significantly more painful than stimuli located distally (low intensity stimuli) in Block 1, Block 2 and Block 3 ‘congruent trials’ (t_(19)_ = 5.80 *p*<0.001, t_(19)_ = 4.03 *p*<0.001, t_(19)_ = 5.30 *p*<0.001, respectively). This was also the case when considering stimuli within each individual axis, except for axis 1 during Block 1 (Figure 3). Results for Bock 3 ‘incongruent trials’ revealed that on average, high and low intensity trials were no longer perceived as significantly different, although significance was still reach when considering axis 3 and axis 7 independently (t_(19)_ = 2.86 *p* = 0.01, t_(19)_ = 2.44 *p* = 0.02, respectively).

**Figure 3.**
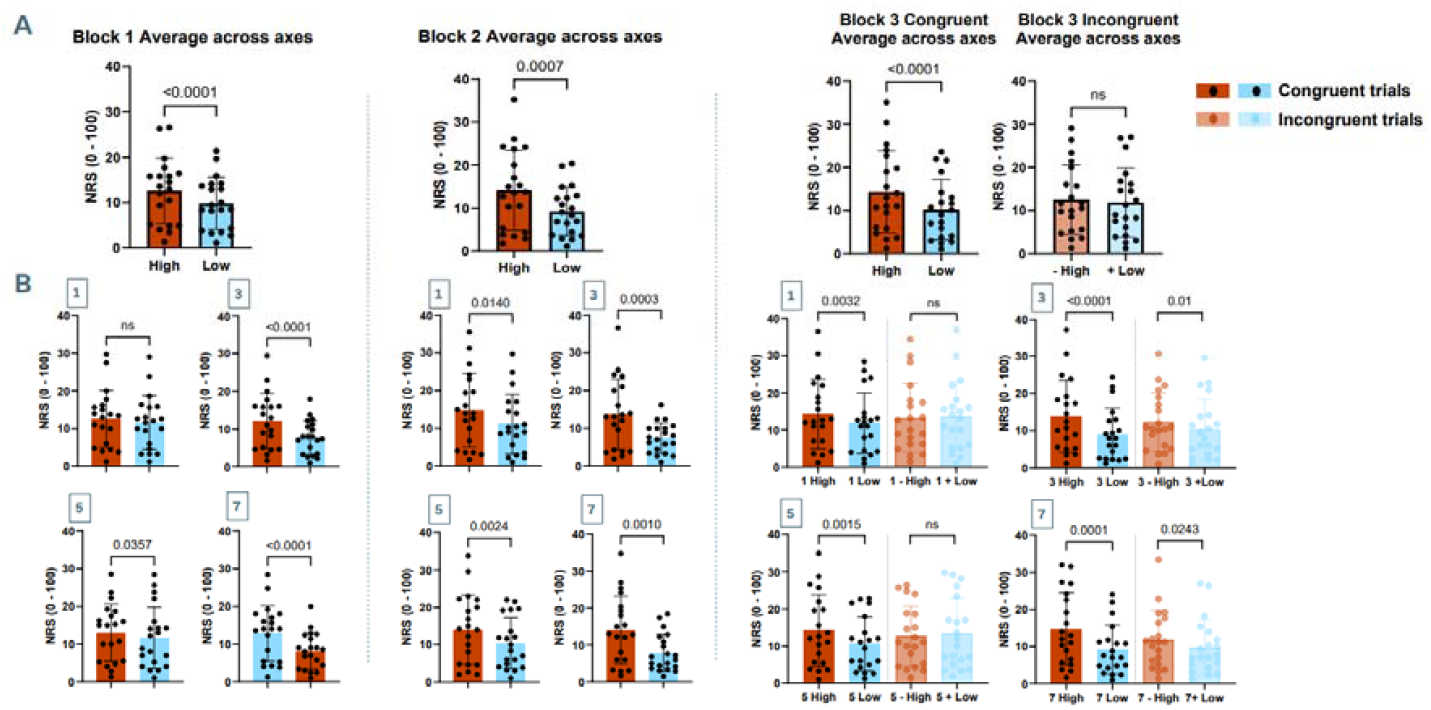
Summary of perceived pain intensity results across blocks for high (red) and low (blue) intensity stimuli. A) perceived pain rating results for high and low intensity stimuli across all axes. Lighter colour bar plots represent results from high and low intensity stimuli preceded by incongruent visual cues. B) perceived pain rating results for high and low intensity trials within each axis. Paired-samples t-tests indicated that high intensity stimuli were perceived as significantly more painful than low intensity trials in most congruent trials, except for axis 1 in Block 1 (second row). There were no significant differences between high and average low intensity incongruent trials for average NRS, axis 1 and axis 5. Error bars represent the SEM. NRS = numerical rating scale (from 0 ‘no pain at all’ to 100 ‘worst pain imaginable’).

### PEs and congruency

One-sample t-tests for PEs within high and low intensity trials for each group (Block 2, Block 3 congruent trials, Block 3 incongruent trials) revealed no significant differences from zero for high and low intensity stimuli in Block 2 and Block 3 congruent trials (Figure 4). PEs were significantly greater than zero for incongruent negative trials (- High), indicating that participants perceived on average significantly more pain than predicted (t_(19)_ = 4.15 *p*<0.001). Conversely, PEs were significant lower than zero for incongruent positive trials (+ Low), reflecting that participants perceived less pain than predicted during these stimuli (t_(19)_ = 4.86 *p*<0.001). One-way ANOVA revealed a main effect of stimulus intensity (F_(2,38)_ = 19.02 *p* <0.001, partial η^2^ = 0.48) with PEs for high intensity trials being significantly greater than low intensity trials across conditions (Mean difference = 1.64 *p*<0.001) and a significant interaction effect (F_(2,38)_ = 17.96 *p* <0.001, partial η^2^ = 0.48), driven by significant PE differences between high and low intensity trials within the Block 3 incongruent block but not in the others (Figure 4).

**Figure 4.**
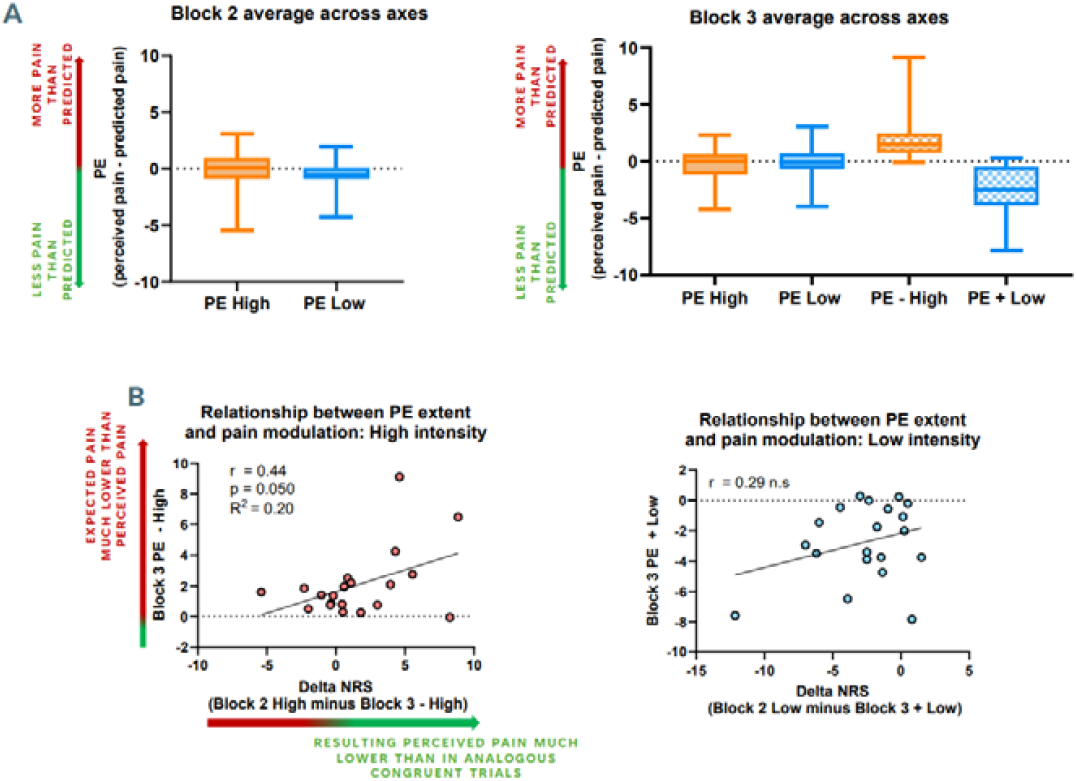
Summary of PE results. A) Group level results for PEs following high (orange) and low (blue) intensity trials during Block 2 (left panel) and Block 3 (right panel). PEs across all congruent trials were not significantly different from zero, indicating high prediction accuracy. PEs for incongruent negative trials were significantly greater than zero, reflecting significantly greater pain than expected, and PEs for incongruent positive trials were significantly lower than zero, reflecting significantly lower pain than expected. B) left panel: Pearson’s correlation between delta scores (resulting from subtracting NRS scores from high intensity trials in Block 2 minus NRS scores from analogous incongruent trials, x axis) and PEs for incongruent negative stimuli (y axis). A significant positive correlation was identified. Right panel: homologous scatter plot for low intensity trials. No significant correlation was found. PE = prediction error; NRS = numerical rating scale; n.s = not significant.

### PEs and pain modulation

We carried out paired-samples t-test to examine whether stimulus congruency influenced perceived pain intensity. There was a significant difference between congruent trials in Block 2 and analogous incongruent trials in Block 3. Concretely, low intensity stimuli in Block 2 (i.e., when a congruent visual cue was provided) were perceived as less painful than in Block 3 incongruent trials (i.e., when an incongruent, high intensity visual cue was provided), t_(19)_ = 3.56 *p = 0*.*002. We* found no significance difference for perception of high intensity stimuli in Block 2 and in Block 3 incongruent trials. Pearson’s correlations revealed that this difference in perceived pain across blocks correlated positively with PEs for incongruent trials of the same stimulus intensity for high intensity trials (Figure 4b). This is, participants who predicted much lower pain than perceived in incongruent negative trials still reported lower pain than in analogous congruent trials, and viceversa. We found no significant correlation for low intensity stimuli.

### Learning effects

While our first exploratory analysis revealed no significant differences between average PEs from the first and second halves of trials of each group (Block 3 congruent, Block 3 incongruent negative, Block 3 incongruent positive), linear regression analyses indicated that there was a significant tendency to zero (i.e., no error of prediction) for PEs in incongruent low intensity trials as a function of trial number (F_(1,18)_ = 6.25 *p* = 0.02, R^2^ = 0.26, Y = 0.046X−5.003). PEs were not significantly predicted by trial number for congruent or incongruent high intensity trials (Figure 5).

**Figure 5.**
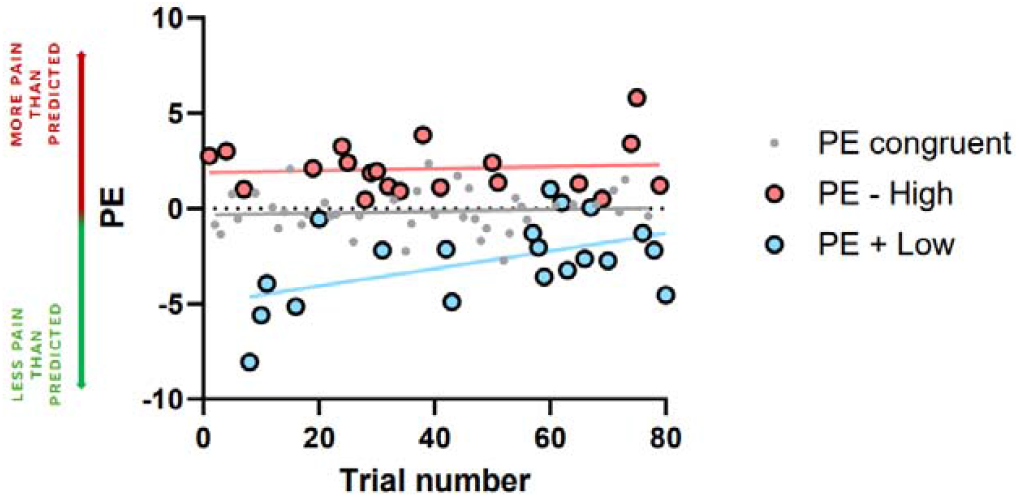
Exploratory linear regression analyses on potential learning effects on PEs in Block 3. PEs significantly tended to zero as a function of trial number in the case of incongruent positive trials (+ Low, blue-coloured dots), but not in the case of incongruent negative (- High, salmon-coloured dots) and congruent trials (grey coloured dots). PE = prediction error.

## Discussion

This study introduces a novel VR-based approach to examine the relationship between PEs and mechanical secondary hyperalgesia within the predictive coding framework of pain in healthy individuals. All preregistered hypotheses were confirmed: HFS successfully induced secondary hyperalgesia, as evidenced by increased MPS in the heterotopic area from HFS stimulation; a priori defined high and low intensity stimulation sites were correctly perceived as more and less painful, respectively; and PEs were effectively generated by manipulating participants’ pain expectations, impacting subsequent pain perception. Crucially, exploratory analyses revealed adaptive updating of PEs (tending toward zero) for low intensity stimuli (i.e., stimuli on non-sensitised areas). In contrast, high intensity stimuli (i.e, stimuli on sensitised areas) showed persistent PEs, resembling maladaptive coding observed in chronic pain. This underscores the potential of this model to assess pharmacological, cognitive, and psychological interventions in predictive coding under controlled sensitised states.

HFS successfully elicited secondary mechanical hyperalgesia, consistent with previous findings employing the same adaptation of the DFNS MPS protocol [40] and similar mechanical assessments [17; 71; 74; 75]. This phenomenon has been described in the literature as a manifestation of long-term potentiation of synapses between primary afferent C-fibres and second level afferent neurons in the superficial dorsal horn of the spinal cord [30; 36; 42; 52; 70], and it is therefore considered a robust model of CS [33; 64; 67; 69; 78]. Since CS is thought to be an important neurophysiological mechanism contributing to the establishment and maintenance of chronic pain, especially in the case of inflammatory and neuropathic pain [19; 22; 76; 81], the HFS model has been used to assess the influence of top-down cognitive components of pain perception during experimentally controlled sensitised states, such as expectations [27; 68], fear and empathy [64], or cognitive demand [63] as well as to provide therapeutics biomarkers [40; 41; 52]. However, to the best of our knowledge, this is the first attempt to characterise patterns of PE coding and pain inference during CS in healthy individuals using this model. Furthermore, we hypothesised that the limited spread of HFS-induced secondary hyperalgesia [40] would allow precise targeting of proximal locations that are differently affected by varying levels of CS; this was largely confirmed by our results, as participant perceived the so called ‘high intensity’ stimuli as significantly more painful than ‘low intensity’ stimuli. Since the volar forearm has a relatively uniform sensory innervation through median and ulnar nerves [7], these results were unlikely to arise due to simple pain sensitivity variations.

Our protocol successfully induced prediction PEs by manipulating the congruency between the shown stimulus location and the actual stimulus location. These PEs occurred in the hypothesised direction: participants perceived greater pain when stimulated at a high intensity landmark while shown a low intensity landmark, and less pain when the reverse was true. Importantly, expectations influenced subsequent pain perception; low intensity stimuli were rated as more painful when participants anticipated high intensity pain, consistent with the principles of nocebo hyperalgesia [8; 10; 24; 65]. In contrast, high intensity stimuli were not rated as significantly less painful when participants anticipated low-intensity pain compared to congruent expectations. This finding diverges from the principles of placebo analgesia [9; 10; 27; 65], where positive expectations are thought to engage endogenous opioid pathways that dampen nociceptive input [5; 15; 47; 80]. It is reasonable to argue that this discrepancy could be attributed to the influence of CS within the peak hyperalgesic area, where amplified nociceptive drive overrides top-down modulatory mechanisms of placebo analgesia [35]. Further, previous evidence suggests that the direction of the mismatch is encoded by differential brain mechanisms [48], which may be differently affected by CS. There was nevertheless a significant correlation between PEs and the extent of placebo analgesia-like effects during high intensity trials, underscoring the critical role of expectations in pain modulation, even in the presence of CS. Conversely, during low intensity stimulation (i.e., where nociceptive input is less amplified), PEs appeared to play a diminished role in pain modulation, consistent with previous findings in thermal pain [59]. This is also reflected in our data by a lack of significant correlation between PEs and pain modulation during low intensity trials. These findings demonstrate how CS may alter the interplay between predictive coding and nociceptive processing, providing a mechanistic explanation for the persistence of maladaptive pain predictions in chronic pain [11; 16]. Follow up studies may explore how trait-like bias, such as pessimistic tendencies during pain prediction [26] or general fear of pain [2; 18; 61], interact with the effects observed in the present study. Such insights have profound implications for understanding pain perception and developing strategies to recalibrate maladaptive predictive coding in sensitised states.

The contrasting patterns of PE adaptation over time observed offer insights into how central CS disrupts predictive coding of pain. When participants were shown a low intensity landmark while poked in a highly sensitised area, PEs remained constant across trials, suggesting that the heightened nociceptive input from sensitised regions may impair the brain’s ability to update its predictions. This rigidity suggests that in CS, the brain prioritises the amplified nociceptive signals over sensory feedback, preventing error correction [9; 43; 51; 56]. Conversely, when participants were shown a landmark in a highly sensitised area while poked in a less sensitised area, PEs diminished across trials, reflecting normal adaptive learning to maximise precision, according to the principles of predictive processing [20; 26; 60] and reinforcement learning [55]. These findings imply that the presence of CS may alter the balance between prior expectations and sensory updates, leading to a maladaptive persistence of predictions in sensitised regions [73]. This phenomenon may mirror mechanisms underlying chronic pain, where excessive nociceptive signals perpetuate inaccurate predictions and hinder the resolution of pain, contributing to its persistence [44; 49].

This model provides a controlled framework to investigate how CS affects predictive coding and pain perception, with potential applications in both research and clinical settings. Moving forward, it could be used to assess how different interventions (pharmacological, psychological, or cognitive) impact predictive coding in the presence of CS [6; 28]. For example, treatments aimed at reducing hyperalgesia or modulating expectations (e.g., cognitive-behavioural therapy, mindfulness-based stress reduction) [29; 66] or promoting placebo analgesia (e.g., hypnosis, placebo interventions) [54] could be evaluated by examining changes PEs and pain modulation patterns. In clinical contexts, this VR model could help identify individuals with maladaptive predictive coding, offering a tailored approach to treatment [58; 79]. Disentangling the roles of nociceptive input and expectation updating could provide insights into maladaptive mechanisms linked to chronic pain but not fully explained by the presence of CS [72]. Additionally, integrating this model into neuroimaging or neurophysiological studies could reveal the neural circuits disrupted by CS and how they respond to therapeutic interventions [12; 43; 45; 51]. Finally, the model could inform the design of VR-based rehabilitation tools [38; 39], using controlled manipulations of visual feedback to retrain the brain’s predictive coding processes and reduce maladaptive pain perceptions.

This study also presents limitations that should be acknowledged. Firstly, the unbalanced female-to-male ratio could have introduced potential gender-related bias regarding pain estimation [83] and pain perception mechanisms during secondary hyperalgesia [1]. Secondly, the absence of a baseline VR task assessment prior to HFS, or of similar assessments in the control arm following HFS, limits our ability to disentangle the specific effects of HFS-induced central sensitisation on predictive coding from participants’ baseline predictive processes. While the focus of this study was to assess the viability of the VR task during HFS, control assessments for the VR task may be implemented in future iterations of this model. Finally, it is worth considering that while participants might have consciously realised the mismatch between the poked location and the visual cue incongruent trials, it may not necessarily be a critical limitation for studying PEs; Predictive Coding Theory posits that PEs arise automatically when sensory inputs deviate from predictions, regardless of conscious awareness [50]. Therefore, even if participants consciously recognise the mismatch, the underlying neural mechanisms driving PEs are likely still engaged. Nonetheless, these considerations highlight the need for further studies to validate and extend our findings.

In summary, this study presents a novel VR-based approach to explore the interplay between PEs and secondary mechanical hyperalgesia in healthy individuals. We successfully demonstrated that expectations influence pain perception in sensitised and non-sensitised areas differently. Importantly, the findings suggest that in sensitised states, PEs persist, potentially reflecting maladaptive predictive coding mechanisms similar to those implicated in chronic pain. Therefore, we propose this model as a potential viable candidate to provide a controlled framework to study interventions targeting maladaptive PE mechanisms. By enabling precise manipulation of expectations and nociceptive input, this new experimental approach can help to inform the development of tailored VR-based rehabilitation strategies targeting PE mechanisms during chronic pain. These findings pave the way for future research to refine the model, validate its clinical relevance, and explore its applications in pain management and rehabilitation.

## Acknowledgements

This study was funded through an Academy of Medical Sciences Springboard grant (SBF007\100108). The author SM is funded by the National Institute for Health and Care Research (NIHR) Exeter Biomedical Research Centre (BRC). The views expressed are those of the author(s) and not necessarily those of the NIHR or the Department of Health and Social Care. The analysis plan for this study (excluding exploratory analyses) was preregistered and available in Open Science Framework (OSF) Registries on the following link https://osf.io/h897k. Preregistration includes study design, variables, and treatment conditions and description of the analysis plan (including specification of the sequence of analyses reported).

